# Radiographic assessment of bone maturation as a tool for age estimation in common dolphins (*Delphinus delphis*)

**DOI:** 10.64898/2026.04.05.716530

**Authors:** Eva-Maria F. Hanninger, Ashley Barratclough, Emma L. Betty, Marti J. Anderson, Matthew R. Perrott, Joy Bowler, Emily I. Palmer, Katharina J. Peters, Karen A. Stockin

**Affiliations:** Cetacean Ecology Research Group, School of Natural Sciences, Massey University, Auckland, New Zealand; National Marine Mammal Foundation, San Diego, United States; New Zealand Institute for Advanced Study, Massey University, Auckland, New Zealand; School of Veterinary Sciences, Massey University, Palmerston North, New Zealand; Auckland Radiology Group, Auckland, New Zealand; Marine Vertebrate Ecology Lab, Environmental Futures, School of Science, University of Wollongong, Wollongong, Australia

**Keywords:** Skeletal maturation, bone ossification, tooth ageing, life-history, cetaceans

## Abstract

We present the first radiographic ageing framework for common dolphins (*Delphinus delphis*), based on ossification and epiphyseal fusion patterns in the pectoral flipper, demonstrating higher reliability for chronological age estimation than currently available epigenetic approaches for this species. Using individuals of known dental age, we calibrated two modelling approaches to predict dental age from radiographic bone scores: 1) a univariate polynomial regression using a total bone score (sum of 16 scores across all assessed flipper bones), and 2) a multivariate canonical analysis of principal coordinates (CAP) incorporating 16 individual bone-score variables. Both approaches successfully predicted dental age from skeletal ossification patterns. For an age range of 0 to 24 years, polynomial regression demonstrated high predictive accuracy with median absolute errors (MAEs) of 1.25 years in females (Spearman’s ρ = 0.93, R² = 0.90) and 1.08 years in males (ρ = 0.95, R² = 0.86). The CAP model yielded MAEs of 1.35 years in females (ρ = 0.90, R² = 0.85) and 1.80 years in males (ρ = 0.94, R² = 0.84). Notably, both radiographic bone ageing models achieved equal or lower median absolute errors and higher coefficients of determination than a recently developed epigenetic clock for common dolphins derived from the same population (MAE = 1.80, Pearson’s correlation (*r*) = 0.91, R² = 0.82). When applying the bone ageing models to individuals of unknown dental age, both models produced age estimates consistent with expected life-history stages (foetus, neonate, juvenile, subadult, adult), although accuracy declined in dolphins above 20 years, likely as a consequence of subtle age-related variation in skeletal changes in this species. Radiographic ageing provides an accurate non-invasive tool for demographic assessment to support conservation management of common dolphins.

## 1. Introduction

Understanding key reproductive parameters, such as age at sexual maturity, gestation and lactation periods, and calving intervals, is central to understanding population viability. Crucially, these assessments rely on accurate age estimation, which provides the foundation for interpreting reproductive status, population structure, and longevity (Betty et al., 2022, 2023). Robust age data are, therefore, essential for evaluating population persistence, predicting responses to environmental and anthropogenic pressures, and informing effective conservation strategies (Betty et al., 2019; Heydenrych et al., 2021; Manlik et al., 2016; Moore & Read, 2008; Palmer et al., 2022).

Traditionally, chronological age assessment in odontocetes has mostly relied on counting growth layer groups (GLGs) in thin, stained tooth sections (e.g., Betty et al., 2023; Evans et al., 2002; Lockyer, 1993, 1995; Murphy et al., 2014; Palmer et al., 2022, 2023; Westgate & Read, 2007). Because tooth extraction is invasive, it is typically confined to post-mortem examinations, with the exception of procedures performed under anaesthesia (Barratclough, Wells, et al., 2019). In addition, GLG assessment requires experienced readers and access to specialised histology laboratories, which can be costly and time-consuming (Evans et al., 2011; Murphy et al., 2014). These logistical and financial constraints further limit the feasibility and scalability of traditional age estimation across populations and regions, highlighting the need for alternative approaches.

Epigenetic age estimation based on DNA methylation provides a feasible non-lethal alternative for ageing live animals (e.g., Barratclough et al., 2021; Hanninger et al., 2025; Hernandez et al., 2023; Peters et al., 2023; Robeck et al., 2021; Zoller et al., 2025), though it also has practical constraints. Implementing these methods requires either high-throughput sequencing or the use of Illumina methylation arrays (Arneson et al., 2022; X. Liu et al., 2024). At present, the relevant patent for array-based processing is held by the Clock Foundation in the United States (Arneson et al., 2022; Lu et al., 2023). Consequently, both the laboratory procedures and the potential need to ship samples internationally are costly and logistically demanding, limiting the accessibility of epigenetic ageing in many research and management contexts.

Such constraints have encouraged interest in methods that are more practical and broadly accessible. Pectoral flipper radiography offers one such option, as radiography is comparatively inexpensive and relies on equipment that is widely available, even in remote field settings. This technique allows age estimation, as recently demonstrated for bottlenose dolphins (*Tursiops truncatus*; Barratclough, Sanz-Requena, et al., 2019), and provides a non-invasive morphological alternative to tooth-based ageing. Based on the assessment of 16 specific flipper bone locations, the method evaluates ossification and epiphyseal fusion, which follow a predictable age-related sequence (Barratclough, Sanz-Requena, et al., 2019). The framework, so far only applied to bottlenose dolphins under human care, incorporates assessments of long-bone maturation, delta-bone development (a cetacean-specific physiological stage in the ossification of specific metacarpals and phalanges; Barratclough, Sanz-Requena, et al., 2019; Dawson, 2003) and the occurrence of age-related degeneration (Barratclough, Sanz-Requena, et al., 2019). However, while the radiographic age assessment framework for bottlenose dolphins represents a major advance, differences in ossification timing between sexes and among species mean that models of ageing must be developed and calibrated on a sex- and species-specific basis, using radiographs from known-aged individuals (Barratclough, Sanz-Requena, et al., 2019).

Species-specific calibration is necessary because developmental trajectories are shaped by life-history traits, including hormonal changes during puberty (Almeida et al., 2017; Iuliano-Burns et al., 2009). Furthermore, the direct applicability of this framework to other cetacean species has remained uncertain, and equivalent models for species such as common dolphins (*Delphinus delphis*) have, until now, been lacking. Uncertainties about transferability arise from morphological differences in flipper anatomy (e.g., degree of hyperphalangy; Cooper et al., 2007), interspecific variation in delta bone maturation, and limited knowledge of the occurrence and extent of degenerative bone disease in other species (Hanninger et al., 2026).

Recently, Hanninger et al. (in press) addressed whether interspecific morphological differences would require modification of scoring criteria for application to common dolphins. While interspecific differences in morphology and bone maturation patterns were present, these differences did not prevent the scoring framework from being applied effectively. Importantly, scoring agreement between independent observers remained high, highlighting the robustness of the method. One potential limitation identified by the authors, however, was that age-related degeneration appeared comparatively more subtle in common dolphins, which could theoretically reduce accuracy in aged individuals (>20 yrs; Hanninger et al., in press).

Building on these findings, the present study develops a calibrated radiographic model for age assessment in common dolphins, providing a species-specific tool for age estimation. The model was developed using a 30-year tissue archive of stranded and bycaught individuals with known dental ages, sex, body length, and sexual maturity (Palmer et al., 2022, 2023). Radiographic bone scores for 137 individuals (aged from 0 to 31 years) were assessed *a priori* (Hanninger et al., 2026) and used to construct a model spanning the full lifespan of free-ranging dolphins (∼30 years; Perrin, 2009) represented in the sample. To account for sex-specific differences in ossification, separate models were developed for males and females. We present two complementary modelling approaches for age estimation. The first approach followed Barratclough, Sanz-Requena, et al. (2019) using the total bone score for each radiograph (i.e., the sum of all individual scores) in a polynomial regression calibrated against dental age. This approach was applied to ensure comparability with previous bottlenose dolphin studies. Second, we used a multivariate approach — canonical analysis of principal coordinates (CAP, Anderson & Robinson, 2003; Anderson & Willis, 2003), where the 16 bone score values were treated as 16 individual inter-correlated variables. In this analysis, principal coordinates are calculated from Euclidean distances among all individuals on the basis of the 16 bone variables, in a holistic framework. The model seeks a linear combination of the principal coordinates to maximise discrimination among dolphin individuals along a gradient of known dental age. This approach allowed the multidimensional structure of flipper ossification to be characterised and related directly to chronological age.

## 2. Materials & Methods

### 2.1. Sample collection

#### 2.1.1. Life-history assessment

Dolphins assessed in this study (*n* = 137) are stranded or bycaught individuals examined postmortem between 2002 and 2025. Age, growth, and sexual maturity were assessed *a priori* to pectoral flipper examination as part of earlier life history research (Palmer et al., 2022, 2023). We included foetuses (*n* = 5; either extracted in utero or aborted prior to first breath, confirmed by pulmonary atelectasis; Colegrove et al., 2016), neonates (*n* = 12; live birth confirmed by inflated lungs and showing at least one of the following characteristics: presence of vibrissal hairs, an unhealed navel, foetal folds, soft and folded dorsal fin and tail flukes, or unerupted teeth; Díaz López et al., 2018; Puig-Lozano et al., 2020), juveniles (*n* = 56; lacking neonatal characteristics and with total body length less than average length at attainment of sexual maturity, i.e. <183 cm for females and <190 cm for males, Palmer et al., 2022, 2023), subadults (*n* = 5; adult-sized, i.e. ≥183 cm in females and ≥190 cm in males, but not yet sexually mature; Palmer et al., 2022, 2023), and adults (*n* = 57; sexually mature). Two additional individuals for which age class could not be determined were also included.

Sex (females: *n* = 72; males: *n* = 65) was determined anatomically during post-mortem examination. In rare cases where the genital region was damaged (*n* = 2), molecular sexing was conducted using multiplex PCR amplification of sex chromosome-specific loci (ZFX and SRY) from skin samples (Gilson et al., 1998; Peters et al., 2023). PCR products were amplified with established primers (P153Z & P23EX and y53-3c & Y53-3D; Aasen & Medrano, 1990; Gilson et al., 1998) and visualised by gel electrophoresis.

Dental age was available for 120 animals (64 females, 56 males). The remaining 17 individuals lacked suitable teeth for GLG ageing (e.g., non-erupted teeth in foetuses and neonates, or teeth otherwise unsuitable for processing). The age was determined by counting GLGs in the dentine of thin, decalcified, and stained tooth sections (Palmer, 2023). Each tooth was independently read by at least two experienced readers across three sessions. If age estimates differed, an additional tooth was processed (up to a maximum of three teeth per individual) and re-read to reach consensus (Palmer, 2023). Dental age ranged from 0 to 31 years, and age distributions by sex are shown in the Supplementary Materials (Figure S1).

#### 2.1.2. Pectoral flipper collection and radiography

Pectoral flippers (*n* = 137 pairs), collected *a priori* during post-mortem examinations, were stored flat, frozen, and subsequently radiographed in a standardized manner, using a single Shimadzu radiographic unit set at 40 kVp (kilovoltage peak) and 4 mAs (milliampere-seconds), with a source-to-sample distance of 100 cm (Hanninger et al., 2026).

### 2.2. Radiographic analysis

Radiographs were evaluated *a priori* in a proximal-to-distal sequence (Hanninger et al., 2026), beginning at the distal radius and ulna and continuing through the metacarpals and phalanges, following Barratclough, Sanz-Requena, et al. (2019). Anatomical bone sites were selected based on their age-informative value in bottlenose dolphins (Barratclough, Sanz-Requena, et al., 2019), as well as their high correlation with age in common dolphins (Hanninger et al., 2026). The scoring locations included the distal epiphyses of radius and ulna, proximal and distal epiphyses of metacarpals II–IV and the first and second phalanges of digits II and III, as well as two delta bones, namely metacarpal V and the first phalanx of digit IV. In contrast, carpal bones, metacarpal I, humerus, and the proximal epiphysis of radius and ulna were excluded, as they either lack clear ossification stages, show weak age associations, or could not be imaged reliably in plane (Barratclough, Sanz-Requena, et al., 2019; Hanninger et al., 2026; Ogden et al., 1981). Scoring protocols were adapted from Barratclough, Sanz-Requena, et al., (2019) and adjusted to reflect morphological variation observed in common dolphins (Hanninger et al., 2026).

Long bones were assigned maturity stages ranging from –1 to 8, reflecting the progression from the initial appearance of ossification centres through physeal narrowing, partial and complete physeal fusion, and, at later stages, degenerative change. Half-scores were applied where intermediate conditions were observed. Long bones were generally scored at both the proximal and distal epiphyses. One exception in the original *Tursiops* framework involved the second phalanges of digits II and III, for which a single combined value was used; this corresponded to the higher of the two epiphyseal scores (Barratclough, Sanz-Requena, et al., 2019). Given that hyperphalangy differs between common dolphins and bottlenose dolphins (Cooper et al., 2007), we evaluated whether this combined-score approach was appropriate for our dataset. To do so, we repeated the analyses under four configurations: treating the proximal and distal scores for these phalanges as separate variables (‘separate epiphyses’), using an averaged combined score (‘combined score average’), applying the maximum score (‘combined score max’), and considering only the minimum score (‘combined score min’).

Delta bones (metacarpal V and the first phalanx of digit four) were scored as single units on a -1 to 8 scale. In common dolphins, both delta bones sometimes exhibited simultaneous development of proximal and distal ossification centres; therefore, for stages 1–4, maturity was classified solely on the proximal ossification centre, regardless of distal development (Hanninger et al., 2026). This approach differs from the scoring system originally developed for bottlenose dolphins, in which delta bone ossification follows a unidirectional sequence, with distal mineralization occurring only after consolidation of the proximal centre (Barratclough, Sanz-Requena, et al., 2019). The classification scheme and criteria used to assign scores to long and delta bones are summarized in Tables 1 and 2, with schematic illustrations available in the Supplementary Materials (Figures S2 and S3). A total bone maturity score was calculated for each radiograph by summing the scores across all assessed locations. Depending on the scoring approach applied to the second phalanx of digits II and III, this total comprised either 16 individual bone scores when using the combined-score method, or 18 individual scores when treating the proximal and distal epiphyses of these phalanges separately.

**Table 1:** Long bone scoring system applied to common dolphins (*Delphinus delphis*) following Barratclough, Sanz-Requena, et al., (2019) with adjustments (*) made based on anatomical differences previously observed for common dolphins (Hanninger et al., 2026).

| Score | Description |
| --- | --- |
| -1 | No primary ossification centre visible |
| 0 | The primary centre is present, but no secondary ossification centre is visible |
| 1 | An epiphyseal ossification centre is present but only occupies <50% of the width of the adjacent metaphysis |
| 2 | The secondary ossification centre is well established, ranging between 50% and 100% of the metaphyseal width; the physis is evident as a distinct radiolucent space, while the secondary centre appears less mature and irregular in depth across its width; maturation progresses in a midline-to-abaxial direction |
| 3 | Thinning of the radiolucent physis but no evidence of fusion of the metaphysis and the secondary ossification centre; mineral density increases in the subchondral bone on both sides of the physis; late stage 2 can resemble early stage 3 with distinction based on whether physis width is irregular (stage 2) or uniform (stage 3) |
| 4 | Osseous bridges begin to form; in the radius and ulna this starts midline and proceeds abaxially; the abaxial margins remain open, and both early bridge formation and >90% bridging qualify as stage 4 |
| 5 | Complete closure is present, accompanied by a faint hyperdense “ghost” physeal line or epiphyseal scar spanning at least 50% of the plate; the former epiphysis may not reach the full transverse width of the bone, but closure is complete |
| 6 | The physeal line is remodelled, with <50% to no evidence of the hypermineralized transverse remnant, and the former epiphysis extends across the full transverse width of the bone; **\*the visibility of the physeal scar was no conclusive criterion in common dolphins and emphasis should be placed on the extension of the former epiphysis** |
| 7 | Complete fusion occurs with no evidence of a previous ghost physeal plate and flattening of the physeal surfaces showing accentuated pointed edges |
| 8 | Arthritic changes are visible, including osteophytes, osteolysis, articular calcification, and in some cases fusion of metacarpals or phalanges |

**Table 2:** Delta bone scoring system applied to common dolphins (*Delphinus delphis*) following Barratclough, Sanz-Requena, et al., (2019) with adjustments (*) made based on anatomical differences previously observed for common dolphins (Hanninger et al., 2026).

| Score | Description |
| --- | --- |
| -1 | Not present |
| 0 | Very small oval shape, undefined delta shape |
| 1 | Axial surface starts to be linear, consistent with delta shape; **\*mineralization may be present adjacent to both the proximal and distal epiphyseal surface but does not show osseous bridge formation** |
| 2 | One flat surface; areas of slight irregular perimetral surface typically on the proximal side with a degree of adjacent mineralization which can oppose the epiphyseal surface but does not necessarily show definitive osseous bridge formation; bone density is clearly reduced in peripheral new bone by comparison to the primary ossification centre; **\*mineralization may be present adjacent to the distal epiphyseal surface and emphasis should be placed on the degree of proximal mineralization** |
| 3 | Increased mineralization of adjacent proximal cartilaginous surface, consolidation may not be present; **\*mineralization may be present adjacent to the distal epiphyseal surface and emphasis should be placed on the degree of proximal mineralization** |
| 4 | Area increases in consolidation and mineralization, slightly less dense than the body of the delta bone; a defined hypermineralized physeal plate line is present; the abaxial aspect can show the initiation of the new straight lateral border rather than a horseshoe shape; **\*mineralization may be present adjacent to the distal epiphyseal surface and emphasis should be placed on the degree of proximal mineralization** |
| 5 | Density of the secondary ossification centre is similar to primary bone and initial mineralization is present on the distal surface, with the proximal side showing a defined straight lateral border rather than a horseshoe shape; individual variation may result in non-linear borders; therefore, emphasis should be placed on the secondary ossification centres on the proximal and distal surfaces |
| 6 | Smooth proximal surface with matching density to the primary ossification centre, distal surface still shows some incongruent mineralization; physeal line is still visible |
| 7 | Smooth proximal surface with complete consolidation of the secondary ossification centre on both sides and dissolution of the physeal line |
| 8 | Degenerative changes present |

### 2.3. Statistical analysis

#### 2.3.1. Bone age modelling

##### i) Polynomial regression

To model the relationship between dental age and skeletal maturity, represented by total radiographic bone score, second-degree polynomial regression models were fitted following Barratclough, Sanz-Requena, et al. (2019). Analyses were conducted separately for females (*n* = 64) and males (*n* = 56) with known dental ages. Predictive performance was evaluated using leave-one-out cross-validation (LOOCV), whereby each individual was excluded in turn, the polynomial regression was refitted to the remaining individuals, and the fitted model was used to predict the excluded individual’s age. This procedure produced one cross-validated age estimate per individual and provided an unbiased assessment of predictive accuracy.

Predicted ages obtained from LOOCV were compared with known dental ages to quantify performance. Accuracy was summarised using the mean, median (MAE), and standard deviation of absolute errors. The association between predicted and dental age was assessed using Spearman’s rank correlation coefficient (ρ) as distributional assumptions for parametric correlation were not met. In addition, linear regression of predicted age on dental age was used to estimate the slope, intercept, and coefficient of determination (R²). Following cross-validation, final polynomial models were refitted using all individuals with known dental ages to derive explicit regression equations, which were subsequently applied to estimate ages for individuals lacking known dental ages. For comparability with the previously published epigenetic clock model (Hanninger et al., 2025), we re-ran the full modelling and cross-validation procedure after excluding individuals with absolute age prediction errors >6 years, as flagged in the polynomial regression and/or CAP model. In the epigenetic analysis, this threshold corresponded to the upper tail of the prediction error distribution and was retained here to ensure methodological consistency across ageing approaches. Performance metrics are reported for both the full and reduced datasets.

Furthermore, as differences in bone maturation between the left and right pectoral flippers were observed in some individuals (∼16%; Hanninger et al., in press), we repeated polynomial regressions for both sexes using five scoring strategies: the lowest total bone score of both flippers (‘BSL’), the highest score (‘BSH’), the average of both flippers (‘ABS’), and the scores from the left (‘Left’) and right (‘Right’) flipper separately. These models were fitted under each configuration of the phalangeal scoring approach (combined scores for the second phalanges of digits II and III using averaged, maximum, or minimum values, or treating proximal and distal epiphyseal scores as individual values), yielding a set of alternative predictive models for each sex.

To assess robustness and practical equivalence across all modelling choices, pairwise comparisons were conducted among all resulting models. For each model pair, absolute differences between cross-validated age predictions (|model A – model B|) were calculated. Agreement between models was summarised using the mean absolute difference and Spearman’s rank correlation coefficient (ρ). Because the distributions of paired differences consistently deviated from normality, no parametric tests were applied. Practical equivalence was evaluated using a nonparametric bootstrap procedure (10,000 resamples) to estimate 90% confidence intervals for the mean absolute difference. Models were considered equivalent when the upper bound of this interval was ≤ 0.5 years. This threshold reflects the typical resolution of dental ageing, which generally cannot reliably distinguish differences smaller than approximately 0.5 years, except in very young individuals where a resolution of ∼0.25 years may occasionally be achievable.

##### ii) Canonical analysis of principal coordinates

Canonical analysis of principal coordinates (CAP; Anderson & Robinson, 2003; Anderson & Willis, 2003) was used to assess whether variation in bone ossification scores across multiple skeletal locations could be used collectively to predict dental age. This approach was chosen because CAP can be used directly to measure and test the extent to which multivariate structure defined by a chosen resemblance measure (in this case, Euclidean distances based on the radiographic bone measurements) can be used to predict positions along a hypothesised variable (in this case, dental age). The canonical model also allows projection of new individuals into the canonical space to predict their positions along the age variable axis. Here, this analysis was applied to determine whether variation in skeletal maturation, captured by 16 (or 18) radiographic bone measurements, can be used to model dental age, and to obtain age predictions for individuals lacking direct age estimates.

All analyses were conducted in PRIMER version 7 (Clarke & Gorley, 2015) with the PERMANOVA+ add-on (Anderson et al., 2008) and were performed separately for females and males. A resemblance matrix was constructed from ossification scores (16 or 18 variables, depending on the scoring approach), with Euclidean distances calculated on untransformed data. Canonical analysis was done on the distance matrix (in each case) to predict dental age. The analysis finds a canonical axis (CAP1) that is a linear combination of principal coordinates (PCO axes, calculated from the distance matrix; Gower, 1966) that achieves a maximum correlation and best discriminates positions along the dental age variable axis. The resulting canonical correlation (δ) captures and quantifies the strength of association between multivariate skeletal development and age.

Not all PCO axes are included in the analysis, but to avoid over-parameterization of the CAP model, a subset of *m* PCO axes is used. The value of *m* is chosen so as to minimize the leave-one-out residual sum of squares of the resulting CAP model (Anderson et al., 2008). Based on this assessment, *m* = 9 was selected for the final models for both females and males; this value captured the majority of age-related structure while avoiding unnecessary model complexity.

Model strength and significance were evaluated using squared canonical correlations (δ²) for the chosen value of *m*, and permutation tests with 9,999 randomisations (Anderson & Robinson, 2003; Anderson & Willis, 2003). A plot of dental age *vs* the CAP axis (drawn separately for males and females) was used to visualise the CAP model. Finally, individuals without known ages were projected onto the CAP axis to obtain their predicted ages, enabling evaluation of their placement along the skeletal maturation trajectory defined by the training dataset.

To evaluate prediction accuracy, CAP-based age estimates were required for all individuals with known dental ages. Leave-one-out cross-validated (LOOCV) age estimates were extracted directly from the CAP analyses and compared with known dental ages to quantify predictive accuracy. Model performance was summarised using the mean, median (MAE), and standard deviation of absolute errors, together with Spearman’s rank correlation coefficient (ρ) between dental and predicted bone ages. In addition, linear regression of predicted age against dental age was used to estimate the slope, intercept, and coefficient of determination (R²). Consistent with the polynomial regression analysis, results are presented for both the full dataset and a subset excluding individuals with absolute prediction errors >6 years in either the polynomial regression or the CAP model, to evaluate robustness to extreme prediction discrepancies.

## 3. Results

### 3.1. Bone age modelling

#### 3.1.1. Polynomial regression

All polynomial regression models showed highly comparable performance across both sexes, with only negligible differences in error metrics and predictive strength across alternative scoring strategies and flipper representations (Supplementary Tables S1 and S2). Furthermore, comparisons of age predictions among flipper representations (‘BSL’, ‘BSH’, ‘ABS’, ‘Left’, ‘Right’) revealed an exceptionally high degree of agreement across all scoring approaches. For females, pairwise comparisons among the 20 models (190 model pairs) showed consistently small differences in predicted ages. Mean absolute errors ranged from 0.02 - 0.13 years (median 0.06 years), and Spearman rank correlations were uniformly high (0.9990 - 1). The upper bounds of the bootstrap 90% confidence intervals for mean absolute error ranged from 0.02 - 0.17 years, well below the 0.5-year practical-equivalence threshold. For males, mean absolute errors ranged from 0.01 - 0.13 years (median 0.06 years), with likewise high Spearman’s rank correlations (0.9976 - 0.9999). The upper bounds of the 90% confidence intervals ranged from 0.02 - 0.18 years, with all model comparisons staying within the 0.5-year threshold. Our results show near-complete consistency in the relative ranking of individuals across models and sexes. All models yielded highly similar age-prediction values, indicating that neither the scoring approach of the second phalanges of digit II and III, nor the occurrence of bilateral differences in bone maturation affected predictive outcomes meaningfully. For subsequent analyses, results are reported for the right flipper using the maximum combined score for the second phalanges of digits II and III. Outputs from all alternative scoring approaches and flipper representations are provided in the Supplementary Materials (Tables S1 to S3).

For females (*n* = 64), the model produced a second-degree polynomial. Age predictions achieved a mean absolute error of 2.03 years, with a median absolute error of 1.36 years and a standard deviation of 2.27 years (Table 3). Predicted ages were strongly correlated with dental ages (Spearman’s ρ = 0.91). Linear regression yielded a slope of 0.82 and an intercept of 1.51 (Figure 1A), with dental age explaining approximately 80% of the variance in predicted age (R² = 0.80). Three individuals exhibited absolute prediction errors >6 years, with a maximum absolute error of 12.77 years. The largest discrepancy occurred in the oldest female (W08-17Dd; 29 years), who appeared radiographically younger relative to her dental age (predicted age: 16.23 years), likely reflecting minimal observable age-related degeneration. In contrast, the other two individuals (absolute errors of 6.28 and 10.26 years) appeared radiographically more mature than expected for their dental age, suggesting advanced skeletal development relative to chronological age. To assess the influence of these pronounced discrepancies on overall model performance, the female model was re-run excluding these individuals (ID codes: W08-17Dd, KS19-17Dd, KS19-20Dd), as well as one additional outlier flagged by the CAP model (KS12-13Dd), resulting in a reduced sample size of 60 females (maximum age 20 years). Under this analysis, mean absolute error decreased to 1.56 years and median absolute error to 1.25 years. The standard deviation of absolute errors was 1.33 years, and the maximum absolute error decreased to 5.99 years. The correlation between predicted and observed ages increased (ρ = 0.93), with an improved regression slope of 0.91, intercept of 0.74, and R² = 0.90 (Figure 1B).

**Table 3:** Error metrics for age predictions in common dolphins (*Delphinus delphis*) generated from sex-specific polynomial models (F = females, M = males). Models were fitted using the total bone score of the right flipper. For each sex, models were additionally recalculated following exclusion of individuals with an age prediction error >6 years in the full model. These outlier-removed versions are denoted as F-WO (females without outliers) and M-WO (males without outliers).

| Error metrics | F | F-WO | M | M-WO |
| --- | --- | --- | --- | --- |
| Mean Absolute Error | 2.03 | 1.56 | 2.13 | 1.71 |
| Median Absolute Error | 1.36 | 1.25 | 1.40 | 1.08 |
| N samples with Absolute Error $> 6$ years | 3 | 0 | 3 | 1 |
| N samples with Absolute Error $> 10$ years | 2 | 0 | 1 | 0 |
| Maximum Absolute Error | 12.77 | 5.99 | 10.95 | 6.23 |

**Figure 1:**
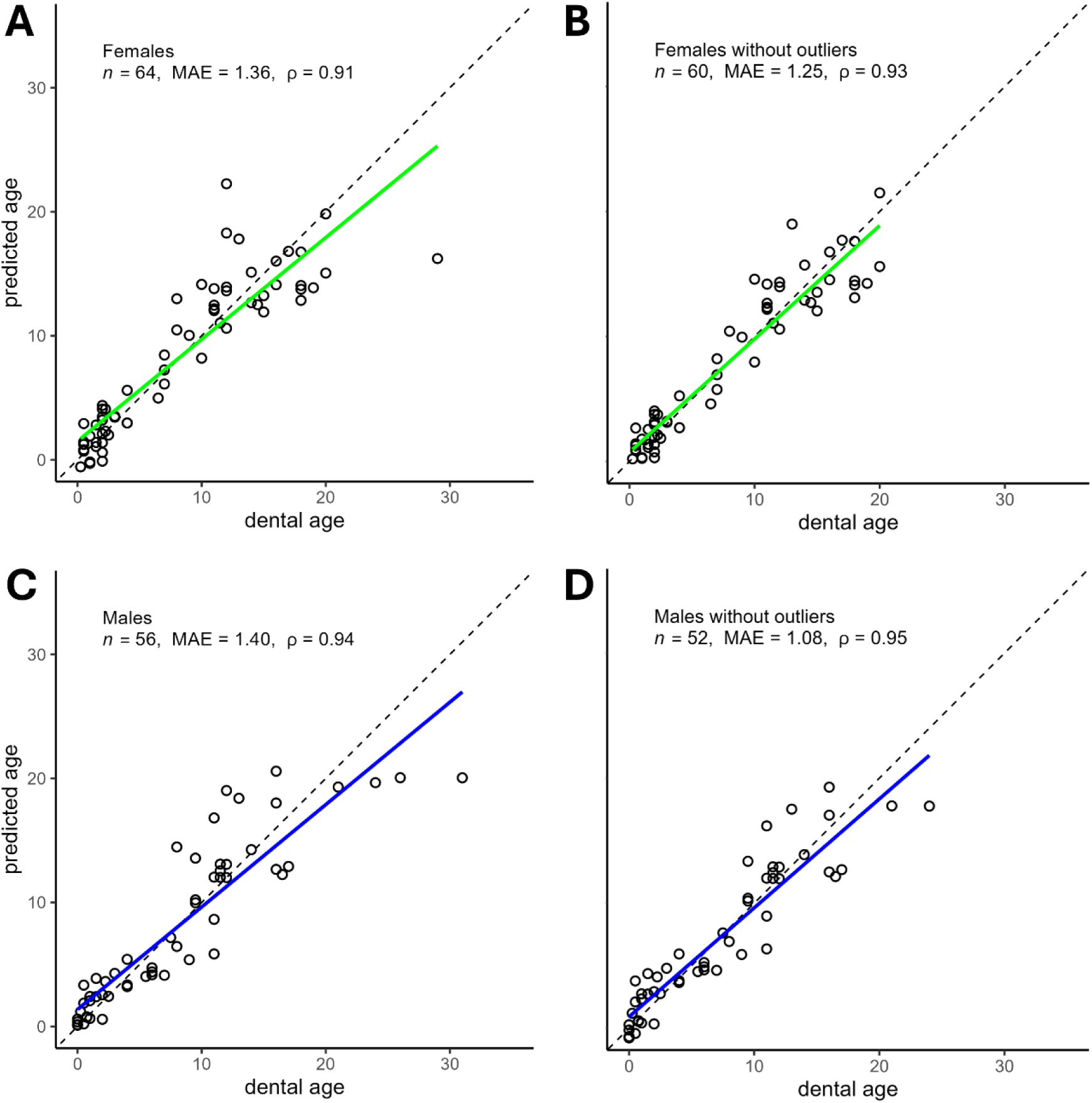
Relationship between dental age and predicted bone age in common dolphins (*Delphinus delphis*). Predicted ages were generated using second-degree polynomial regression of dental age on the total bone score of the right flipper, with leave-one-out cross-validation (LOOCV). A) Model based on the total bone score of the right flipper in females. B) Model based on the total bone score of the right flipper in females but excluding outliers with absolute age prediction errors >6 years. C) Model based on the total bone score of the right flipper in males. D) Model based on the total bone score of the right flipper in males but under exclusion of outliers with absolute age prediction errors >6 years. Each circle represents an individual. Solid-coloured lines show linear regressions between dental age and LOOCV-predicted age, while dashed lines represent the 1:1 line. Reported are sample size (*n*), median absolute error (MAE), and the Spearman rank correlation coefficient (ρ).

For males (*n* = 56), the model produced a second-degree polynomial. Predictions achieved a mean absolute error of 2.13 years, with a median of 1.40 years and a standard deviation of 2.21 years. Predicted ages were very strongly associated with dental age (Spearman’s ρ = 0.94). Linear regression yielded a slope of 0.83 and an intercept of 1.38 (Figure 1C), with dental age explaining about 82% of the variance in predicted age (R² = 0.82). Three males exhibited absolute prediction errors >6 years, with a maximum absolute error of 10.95 years observed in the oldest male (KS23-45Dd; 31 years), who appeared radiographically younger relative to his dental age (predicted age: 20.05 years), likely reflecting limited observable age-related degeneration. The remaining two males showed comparatively advanced skeletal maturation relative to their dental age, with absolute prediction errors of 6.48 and 7.02 years. To assess the influence of these pronounced discrepancies on overall model performance, the model was re-run excluding these individuals (ID codes: KS09-13Dd, KS23-45Dd, KS23-50Dd), as well as one additional animal flagged by the CAP analysis (KS15-16Dd), reducing the sample size to 52 males (maximum age 24 years). This sensitivity analysis resulted in a second-degree polynomial (malesWO) with a mean absolute error of 1.71 years, a median absolute error of 1.08 years, and a standard deviation of absolute errors of 1.52 years. The maximum absolute error decreased to 6.23 years (observed in the oldest remaining male, KS23-46Dd; 24 years). The correlation between predicted and observed ages remained high (ρ = 0.95), with a regression intercept of 0.91, slope of 0.87, and R² = 0.86 (Figure 1D). Error metrics are summarised in Table 3. The corresponding polynomial equations are presented below, where y denotes predicted age (years) and x the total bone score.

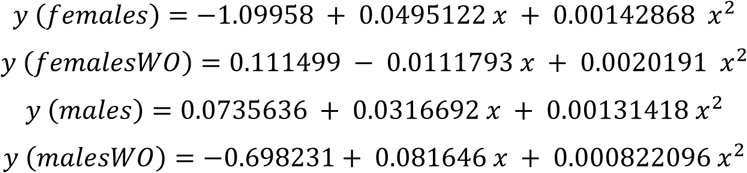

Equations for the other model configurations are provided in the Supplementary Materials (Table S3). Age predictions for animals with unknown chronological ages, estimated using these formulas, are provided in Table 4.

**Table 4:** Estimated ages for common dolphins (*Delphinus delphis*) with unknown or unavailable dental ages, derived using two independent approaches. Ages were predicted using (i) a sex-specific second-degree polynomial regression model based on total pectoral-flipper ossification scores of the right pectoral flipper, and (ii) a Canonical Analysis of Principal Coordinates (CAP) model. Individuals included foetuses, neonates, juveniles, subadults, and adults for which no validated dental ages (GLGs) were available. Both models were fitted on the full dataset. Negative age estimates reflect extrapolation at very early developmental stages where epiphyseal fusion scores fall outside the calibration range.

| ID | Age class | Polynomial age | CAP age |
| --- | --- | --- | --- |
| WS02-44Dd | ND | 0.07 | -1.73 |
| WS07-01DdA | Foetus | -0.58 | -1.20 |
| KS10-78Dd | Subadult | 8.95 | 8.49 |
| KS11-50DdA | Foetus | -1.10 | -2.53 |
| KS13-02Dd | Neonate | 0.24 | -1.54 |
| KS14-62Dd | Neonate | 0.06 | -2.85 |
| KS19-23Dd | Adult | 20.91 | 19.98 |
| KS21-01DdA | Foetus | 0.07 | -1.73 |
| KS22-03DdA | Foetus | -1.10 | -2.53 |
| KS22-73Dd | Foetus | 0.77 | 0.28 |
| KS22-77Dd | Neonate | 0.44 | -2.07 |
| KS24-68Dd | Neonate | -0.26 | -1.00 |
| KS24-88Dd | Juvenile | 3.59 | 3.40 |
| KS24-96Dd | Neonate | 0.07 | -1.73 |
| KS25-11Dd | Neonate | 0.39 | -2.23 |
| KS25-79Dd | Adult | 14.39 | 14.02 |
| KS25-80Dd | Adult | 11.45 | 10.64 |

#### 3.1.2. Canonical analysis of principal coordinates (CAP)

Canonical analysis of principal coordinates (CAP) revealed a strong association between bone ossification scores and dental age. Results are presented for the right pectoral flipper, using the maximum scores for the proximal and distal epiphyses of the second phalanges of digits II and III.

Using the full dataset, the first canonical axis (CAP1) showed a canonical correlation of δ = 0.93 in both sexes, corresponding to a squared canonical correlation (δ₁²) of 0.87, indicating that CAP1, drawn from the multivariate skeletal maturation patterns, explained approximately 87% of the variation in dental age (Figure 2A and B).

**Figure 2.**
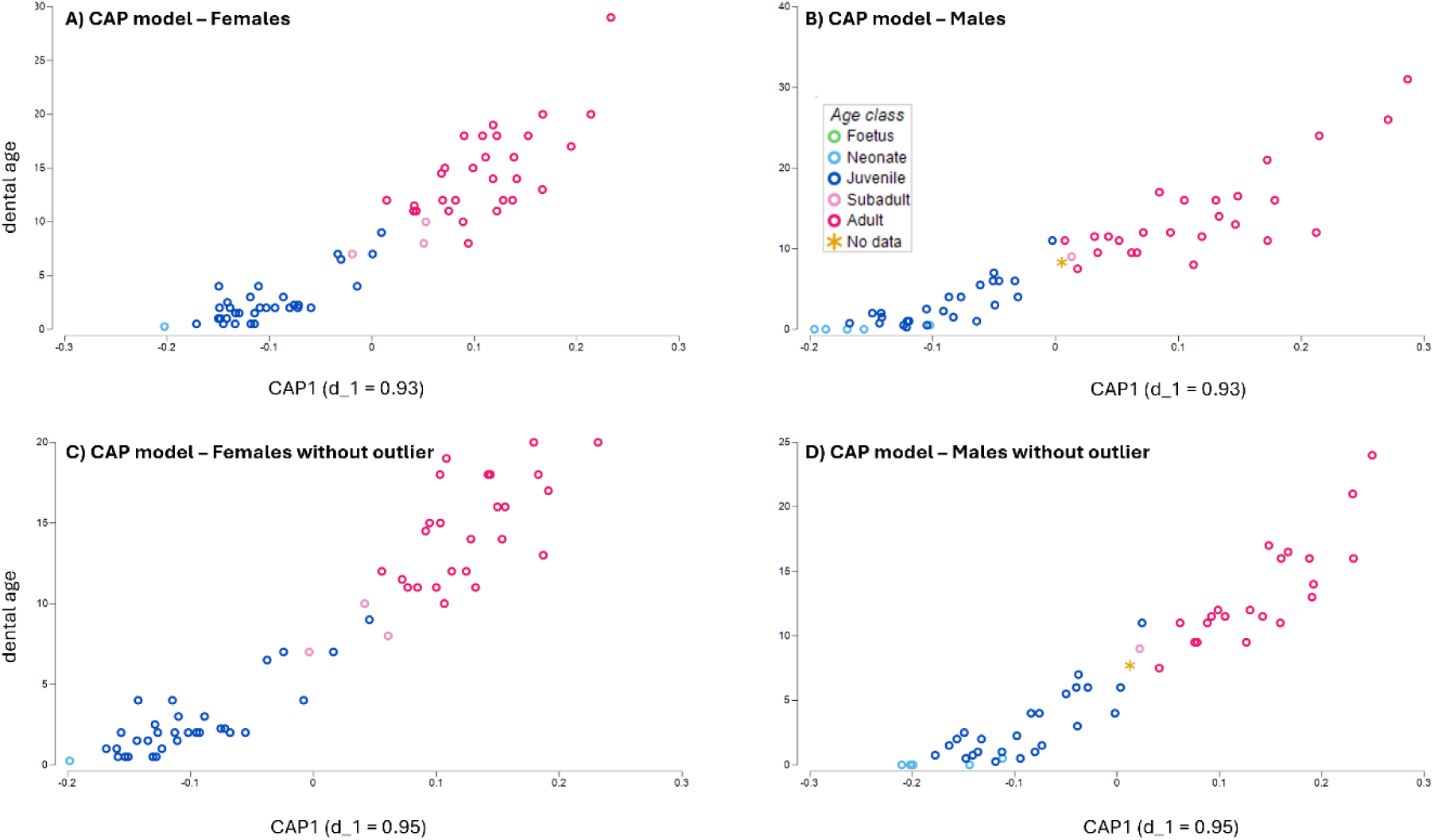
Canonical Analysis of Principal Coordinates (CAP) based on Euclidean distances calculated from 16 pectoral-flipper ossification variables of individual common dolphins (*Delphinus delphis*), showing their multivariate relationship with dental age (in years). Panels show CAP models for A) females and B) males fitted to the full calibration dataset, and C) females and D) males fitted after exclusion of the identified age outliers. In each panel, points represent individuals, coloured according to their assigned age class, as indicated (foetus, neonate, juvenile, subadult, adult, and unknown). CAP1 represents the first canonical axis; d_₁ denotes the canonical correlation between CAP1 and dental age.

CAP-derived fitted values for individuals of known ages showed a mean absolute error of 2.23 years in females (median = 1.79 years; SD = 1.94 years), with a Spearman correlation of ρ = 0.90. Linear regression yielded a slope of 0.85, an intercept of 1.27, and an R² of 0.81. Two females (W08-17Dd, KS12-13Dd) exhibited age prediction errors >6 years. The maximum error was 10.84 years in females. Following exclusion of individuals with age prediction errors >6 years (as identified by either the polynomial regression or CAP model), model performance improved substantially. For females, the canonical correlation increased to 0.95, corresponding to a squared canonical correlation (δ₁²) of 0.90 (Figure 2C). The mean absolute error between dental and predicted age was 1.90 years (SD = 1.60), with a median absolute error of 1.35 years. Spearman’s rank correlation was ρ = 0.90. Linear regression of predicted age on dental age yielded a slope of 0.89, an intercept of 0.86, and R² = 0.85.

In males, the full dataset yielded a mean absolute error of 2.51 years (median = 1.85 years; SD = 2.71), with a Spearman correlation of ρ = 0.93. Linear regression of predicted age on dental age produced a slope of 0.76, an intercept of 1.64, and R² = 0.74. Age prediction error exceeded 6 years in four males (KS09-13Dd, KS15-16Dd, KS23-45Dd, KS23-50Dd), with a maximum error of 13.43 years. Following exclusion of these individuals, the canonical correlation increased to 0.95 (δ₁² = 0.90; Figure 2D). The mean absolute error decreased to 1.90 years (SD = 1.55), with a median absolute error of 1.80 years. Spearman’s ρ was 0.94, and linear regression yielded a slope of 0.88, an intercept of 0.82, and R² = 0.84. Error metrics are summarised in Table 5.

**Table 5:** Error metrics for age predictions in common dolphins (*Delphinus delphis*) generated using sex-specific CAP models (F = females, M = males). Models were fitted using the total bone score of the right flipper. For each sex, models were additionally recalculated following exclusion of individuals with an age prediction error >6 years in the full model. These outlier-removed versions are denoted as F-WO (females without outliers) and M-WO (males without outliers).

| Error metrics | F | F-WO | M | M-WO |
| --- | --- | --- | --- | --- |
| Mean Absolute Error | 2.23 | 1.90 | 2.51 | 1.90 |
| Median Absolute Error | 1.79 | 1.35 | 1.85 | 1.80 |
| N samples with Absolute Error > 6 years | 2 | 1 | 4 | 1 |
| N samples with Absolute Error > 10 years | 1 | 0 | 2 | 0 |
| Maximum Absolute Error | 10.84 | 8.20 | 13.43 | 7.60 |

Finally, the CAP model was applied to individuals of unknown dental ages (Table 4). Foetal and neonatal specimens plotted below the neonatal range, and linear back-prediction from CAP1 produced negative age values.

## 4. Discussion

### 4.1. Advantages of radiographic ageing

This study represents the first radiographic ageing method for common dolphins. Skeletal development of the pectoral flipper can be used to accurately estimate chronological age in this species. Specifically, the strong agreement between radiographically predicted ages and dental ages indicates that bone ageing provides a robust and reliable approach to age estimation in common dolphins. Indeed, both polynomial regression (females: MAE = 1.25, ρ = 0.93, R² = 0.90; males: MAE = 1.08, ρ = 0.95, R² = 0.86) and CAP models (females: MAE = 1.35, Spearman’s ρ = 0.90, R² = 0.85; males: MAE = 1.80, ρ = 0.94, R² = 0.84) demonstrated comparable or improved predictive performance relative to a recently developed epigenetic clock derived from the same population (MAE = 1.80, r = 0.91, R² = 0.82; Hanninger et al., 2025). Furthermore, reliable age estimates up to approximately 20 years of age for both sexes were achieved with bone ageing, representing a substantial advance by comparison with epigenetic approaches, which demonstrated increasing prediction error for animals from ∼16 years of age within the same population (Hanninger et al., 2025).

We acknowledge that bone-based models were fitted separately for females and males, whereas the epigenetic clock was developed using a pooled-sex framework. This structural difference should be considered when interpreting comparative performance. However, empirical evidence suggests that sex-specific stratification of DNA methylation clocks do not generally result in meaningful improvements in predictive accuracy (Carlsen et al., 2023; Sala et al., 2024). To address this structural difference, additional harmonised analyses were conducted. The epigenetic clock published by Hanninger et al. (2025) was refitted independently for females and males, and a pooled polynomial regression model combining both sexes was fitted for the skeletal ageing approach (Supplementary Materials, Text S1–S3, Figures S4–S5). These analyses did not alter the comparative outcome: the skeletal ageing models continued to demonstrate lower absolute prediction error and stronger correlation than the methylation-based model. The superior performance of the radiographic bone ageing method, therefore, cannot be attributed solely to differences in sex stratification between modelling frameworks.

Instead, these differences might reflect the fact that these two methods extract fundamentally different types of information associated with the biology of an ageing organism. Radiographic models capture skeletal growth and maturation, which follow a relatively predictable developmental trajectory closely aligned with chronological age. In contrast, epigenetic clocks quantify age-related molecular changes that reflect biological age and are more sensitive to individual condition, health status, and life history (Barratclough et al., 2024). Although both skeletal development and epigenetic ageing can be influenced by environmental and physiological factors (e.g., Colich et al., 2020; Fitzgerald et al., 2021; Liu et al., 2021; Shirazi et al., 2020; Weindruch et al., 2001), epigenetic age is inherently more sensitive to cumulative stressors and variation in life experience, which can reduce concordance with chronological age (Jylhävä et al., 2017; Levine et al., 2018).

In addition to its demonstrated reliability, radiographic bone ageing offers substantial practical advantages for age estimation in common dolphins. Traditional tooth-based ageing requires tooth extraction followed by laboratory processing involving significant expertise, procedures that are time-consuming, costly, and not always feasible in field-based or resource-limited settings (Betty et al., 2023; Evans et al., 2002; Lockyer, 1993, 1995; Murphy et al., 2014; Palmer et al., 2022, 2023; Westgate & Read, 2007). Moreover, interpretation of growth layer groups becomes increasingly challenging in older individuals, where tooth wear and the accumulation of accessory lines reduce clarity and reliability (Barratclough et al., 2023). Radiographic ageing avoids these constraints and has been shown to provide highly reliable age estimates in bottlenose dolphins, with improved performance relative to GLG-based assessments, particularly at older ages (Barratclough et al., 2023).

Radiographic bone ageing also offers practical advantages when considered alongside molecular age-estimation approaches. While epigenetic ageing enables minimally invasive age estimation in live animals, it depends on specialised and in some cases patented, molecular methods, resulting in substantial per-sample costs (e.g., Hanninger et al., 2025; Hernandez et al., 2023; Peters et al., 2023; Robeck et al., 2021; Zoller et al., 2025). By contrast, radiographs are relatively inexpensive, widely accessible through mainstream human medical facilities, and can be collected routinely in many post-mortem workflows. While on-site imaging during stranding events may not always be feasible, in cases where it is, digital radiographs can be examined in real time, providing immediate age estimates that are highly valuable during live health assessments. Together, these attributes support radiographic ageing as a powerful and cost-effective approach within the broader suite of current age-estimation methods.

### 4.2. Performance of the modelling approaches

To translate radiographic data into age estimates, we applied two modelling approaches. The first was a polynomial regression, adapted from the bottlenose dolphin framework (Barratclough, Sanz-Requena, et al., 2019). This approach showed a high level of stability, even though common dolphins differ anatomically from bottlenose dolphins in ways that could have influenced scoring, including a broader range of hyperphalangy (Cooper et al., 2007) and known lateralised differences in bone maturation (Hanninger et al., 2026). We evaluated whether alternative assessments of the second phalanx of digits II and III, as well as different configurations for combining scores from both flippers, would influence predicted ages. Across all of these variations, the predicted ages remained essentially unchanged. Taken together, these findings indicate that the polynomial approach is robust to minor scoring variation, supporting the use of unilateral radiographs in live animals, where minimising handling time and radiation exposure is a priority. Accordingly, we presented results for the right flipper only using the maximum combined score for the second phalanges of digits II and III, consistent with the previously published bottlenose dolphin framework (Barratclough, Sanz-Requena, et al., 2019). The second modelling approach, CAP (Anderson & Robinson, 2003; Anderson & Willis, 2003), provided a multivariate view of skeletal maturation capturing coordinated developmental patterns. Both approaches captured age-related variation in flipper development and demonstrated high and broadly comparable predictive performance. Together, these results demonstrate that both univariate (polynomial regression) and multivariate (CAP) approaches can be successfully used to estimate age from bone maturation in common dolphins.

The marginal differences in predictive performance between polynomial regression and CAP may be explained by differences in how the two approaches incorporate bone score information. Radiographic bone scoring showed good interobserver consistency in this study (Hanninger et al., 2026), although natural morphological variation can lead to small differences in interpretation, particularly in transitional stages where ossification has not yet fully progressed into a defined maturation category. Polynomial regression aggregates information into a single total bone score by summing across all assessed sites, thereby reducing the influence of minor discrepancies at individual bones and limiting the impact of small scoring differences. In contrast, CAP treats each bone score as an independent variable within a multivariate framework, meaning that small inconsistencies at individual sites may introduce additional noise into the ordination structure. In addition, we used a linear CAP model, whereas the polynomial regression (including a squared term) clearly permits some non-linearity. Thus, this multivariate approach, capturing the holistic shapes in skeletal features across multiple measurements, may well be further improved by the use of a suitable nonlinear CAP model (e.g., Anderson et al., 2005; Millar et al., 2005)

For both the polynomial regression and CAP models, linear regression of predicted age against dental age yielded slopes <1 (polynomial regression: full dataset: females: 0.82, males: 0.83; without outliers: females: 0.91, males: 0.87; CAP: full dataset: females: 0.85, males: 0.76; without outliers: females: 0.89, males: 0.88), indicating compression of the predicted age range and reduced accuracy at older ages. Intercepts were positive across all models. For the polynomial regression, intercepts were 1.51 and 1.38 years in the full dataset and decreased to 0.74 and 0.91 years after outlier exclusion in females and males, respectively. For the CAP models, intercepts were 1.27 years in females and 1.64 years in males for the full dataset. Following exclusion of individuals with absolute prediction errors >6 years, intercepts decreased to 0.86 years in females and 0.82 years in males. In both modelling approaches, removal of extreme prediction discrepancies resulted in slopes shifting closer to 1 and intercepts closer to 0, indicating reduced systematic bias in the linear fit.

Patterns observed at older ages are a direct consequence of this regression behaviour interacting with species-specific skeletal ageing processes. Slopes below 1 result in age-range compression, such that predicted age increases more slowly than true age, leading to age underestimation. In the present study, this pattern appears to be driven by two factors. First, the sample contained relatively few animals above the age of attainment of physical maturity (females: n = 8; males: n = 4; ≥18 years for females and ≥20 years for males; Palmer, 2023). As the study relied entirely on opportunistic strandings, the limited number of confirmed older individuals was likely insufficient to offset regression-to-the-mean effects (Campbell & Kenny, 1999), reducing accuracy at the upper end of the age range.

Second, once physical maturity is reached (≥18 years for females, ≥20 years for males; Palmer, 2023), ongoing skeletal growth no longer provides a strong age-informative signal, and age estimation increasingly relies on age-related degenerative changes. These degenerative skeletal modifications appeared to be far more subtle in common dolphins than in bottlenose dolphins (Hanninger et al., 2026). This may partly reflect differences in the mobility of the animals examined, given that the present study analysed free-ranging common dolphins, whereas the bottlenose dolphin study examined individuals under human care (Barratclough, Sanz-Requena, et al., 2019). Differences observed may also reflect species-specific life-history traits, including longer life expectancy in bottlenose dolphins and differences in the timing of sexual maturity relative to maximum lifespan (Hanninger et al., 2026), which influence growth plate closure. Consequently, while the models of ageing examined here performed robustly across early and mid-life stages, age estimates likely become progressively more variable in geriatric individuals. Nevertheless, our models provided reliable age estimates for both sexes, with substantial age underestimation occurring only in animals older than 20 years.

### 4.3. Radiographic age estimation in foetuses and neonates

Most individuals lacking dental ages in our study were foetal or neonatal specimens with unerupted teeth, precluding tooth-based age determination. As foetal specimens occupy maturation states below those observed in postnatal individuals, both models assigned negative age estimates to foetuses, reflecting their position along a maturation gradient calibrated using postnatal individuals.

Both modelling approaches showed limited ability to distinguish late-term foetuses from very young neonates prior to tooth eruption. The polynomial regression correctly classified three of five foetuses as negatively aged, whereas the CAP model classified four of five foetuses as negatively aged. For neonates, one of six individuals was incorrectly classified as negatively aged by the polynomial regression, and six were classified as negatively aged by the CAP model. When extrapolated beyond the calibrated postnatal age range, both modelling frameworks position these individuals along a similar segment of the maturation gradient, limiting their ability to reliably discriminate between foetal and neonatal stages. Importantly, not all derived polynomial equations can yield negative predicted ages; some formulas remain strictly non-negative across the observed range of bone scores and therefore support only postnatal age estimation.

Beyond statistical reasons for the lack of discrimination, radiographic assessment revealed substantial overlap in skeletal maturation between foetuses and neonates, which contrasts with findings from bottlenose dolphins, where foetuses and neonates could be distinguished radiographically (Barratclough, Sanz-Requena, et al., 2019). However, such discrimination in *Tursiops* may have been driven by the inclusion of very early-stage foetuses (aborted approximately five to six months prior to parturition), whose skeletal development lay well below the neonatal range. In contrast, the present study assessed common dolphin foetuses predominantly with total body lengths above 90 cm, within the reported range of birth lengths for this species (82–101 cm) and close to the estimated mean length at birth (98.1 cm; Palmer, 2023). Consequently, skeletal development in late-term foetuses likely overlaps substantially with that of neonates, reducing the capacity to distinguish these categories radiographically.

Importantly, variation in ossification may not solely reflect gestational stage but may also arise from inter-individual differences in growth trajectories. Notably, the bottlenose dolphin study was based on animals in human care (Barratclough, Sanz-Requena, et al., 2019), where maternal condition, nutrition, and overall health are monitored and likely more consistent. In free-ranging common dolphins, a broader range of maternal factors may influence skeletal development in both foetuses and neonates, potentially accelerating or delaying bone maturation and increasing overlap in radiographic appearance. Such factors include maternal nutrition (Lewall & Cowan, 1963; Ratib et al., 2012; Thacher et al., 2000), underlying disease (e.g., *Brucella ceti* infection; Roca-Monge et al., 2022), and exposure to endocrine-disrupting environmental contaminants (Schwacke et al., 2012) known to adversely affect bone growth (Brankovič et al., 2020; Shulhai et al., 2024; Yaglova & Yaglov, 2021). These environmental and health-related influences may therefore have contributed to the lack of differentiation observed in radiographic age estimates at the earliest life stages. Future research could aim to investigate how late into foetal development foetuses and neonates can be reliably distinguished.

### 4.4. Conclusions

Radiographic scoring of the pectoral flipper is a reliable method for estimating age in common dolphins. This approach is quick, practical, accessible, and provides a comparatively low-cost alternative to tooth-based or molecular ageing methods. The ability to apply this method retrospectively to museums and archives is an additional advantage. Models developed here yielded reliable age estimates for individuals up to approximately 20 years of age in both sexes, with systematic underestimation occurring primarily in individuals older than 20 years. This represents a substantial advance, particularly given that epigenetic ageing approaches in this species show increasing prediction error from approximately 16 years of age onward. Together, these findings underscore the enduring value of morphological and developmental assessments for age estimation and demonstrate that radiographic assessment is a robust and informative tool for life-history research in common dolphins.

At present, this study represents only the second radiographic age-estimation framework developed for cetaceans and the first to explore epigenetic aging of the same individuals comparatively. Our findings highlight the broader relevance of radiographic ageing approaches for species in which other ageing methods are limited or unavailable and emphasise the importance of long-term, multi-tissue archives for advancing life-history research across cetaceans.

## 6. Ethics declaration

Ethic approval was not required as the study was conducted post-mortem on stranded / bycaught animals.

## 8. Data availability

The R code used in this study is openly available on GitHub at https://github.com/Ehanninger/Radiographic-assessment-of-pectoral-flippers. The underlying dataset will be released on GitHub upon publication. The dataset will additionally be released through PRIMER-e as a training resource.

## 9. Author contribution

EMFH contributed to conceptualization, data curation, formal analysis, funding acquisition, investigation, methodology, and draft writing. AB contributed to conceptualization and methodology and supported formal analysis and draft writing. ELB contributed to methodology and supported manuscript drafting. EIP contributed to methodology and supported manuscript drafting. MJA contributed to formal statistical analyses and supported manuscript drafting. MRP supported methodology and manuscript drafting. JB contributed to data curation and methodology. KJP contributed to formal analysis, draft writing, and supervision. KAS contributed to conceptualization, data curation, funding acquisition, methodology, draft writing, and supervision.

## Acknowledgments

Sample collection was carried out under permits 39239-MAR and 111522-MAR issued by the New Zealand Department of Conservation Te Papa Atawhai. We extend our sincere thanks to our iwi partners for granting approval to image their taonga as part of this research. We also acknowledge Alex Burton, Odette Howarth, and the wider Cetacean Ecology Research Group (School of Natural Sciences) for their contributions to sample acquisition, flipper dissection, and dental ageing, as well as Evelyn Lupton and Petru Daniels (School of Veterinary Science) for preparing histological sections of teeth and bones. We further thank Dale Euinton for his support with coding, Pam Aitken and the Auckland Radiology Group team for their support with pectoral flipper imaging, Veronica Candejas for creating the schematic illustrations of long and delta bone development and Emma Carroll (University of Auckland) for genetic sex determination.

