## Supplementary Materials for "Radiographic assessment of bone maturation as a tool for age estimation in common dolphins (*Delphinus delphis*)"

#### Supplementary text

##### Text S1 Extended methods

To ensure a transparent and structurally consistent comparison between skeletal and methylation-based age estimation approaches, additional model variants were fitted. The originally published epigenetic clock (Hanninger et al., 2025) was developed using a pooled-sex framework, whereas the primary polynomial bone ageing models were fitted separately for females and males. To account for this structural difference and avoid confounding model comparison by sex stratification, we implemented two complementary analyses.

First, the epigenetic clock was refitted independently for females and males using the '*restricted*' subset (excluding individuals with an age prediction error >6 years) and analytical procedures described in Hanninger et al. (2025). Second, a pooled-sex polynomial regression model was fitted to the combined male and female dataset following the same procedures outlined in the main methods for skeletal ageing (excluding individuals with an age prediction error > 6 years). These additional analyses were conducted solely to harmonise model structure and to evaluate whether differences in predictive performance could be attributed to sex stratification rather than to the underlying ageing approach.

#### Sex-specific refitting of epigenetic clock published by Hanninger et al. (2005)

To evaluate whether sex-specific stratification influences predictive performance, the epigenetic clock originally described in Hanninger et al. (2025) was refitted independently for females and males using the '*restricted*' subset (excluding individuals with an age prediction error > 6 years). The statistical framework, including elastic net regression and leave-one-out cross-validation, followed the procedures described in Hanninger et al. (2025).

The sole structural modification was that model fitting was conducted independently within each sex rather than on the pooled dataset. In contrast to the original pooled model (Hanninger et al., 2025), which applied a single mean age at attainment of sexual maturity (ASM) across both sexes (8.15 years), the sex-specific refitting incorporated sex-specific average ASM values (female ASM: 7.5 years; male ASM: 8.8 years; Palmer et al., 2022, 2023).

#### Bone ageing models fitted to combined sexes

For completeness, both the polynomial regression, as well as the CAP model were also conducted on the combined male and female dataset. The modelling approach followed the procedures outlined in the main methods section. Outlier exclusion criteria were identical to those applied in the sex-specific analyses; individuals with an age prediction error > 6 years in the model fitted to the full dataset were excluded. No additional model modifications were implemented.

### Text S2 Extended results

#### Sex-specific refitting of epigenetic clock published by Hanninger et al. (2005)

Sex-specific refitting of the epigenetic clock in females ($n$ = 36) yielded a Pearson correlation of $r$ = 0.89 ($p = 1.95 \times 10^{-13}$) and an $R^2$ of 0.80 (Figure S4A). The regression slope of predicted on observed age was 0.62 with an intercept of 2.63 years. Mean absolute error was 2.72 years, and the median absolute error (MAE) was 2.31 years. Two individuals exhibited absolute prediction errors exceeding 6 years, with a maximum absolute error of 8.8 years.

Sex-specific refitting of the epigenetic clock in males ($n$ = 27) yielded a Pearson correlation of $r$ = 0.80 ($p = 5.45 \times 10^{-7}$) and an $R^2$ of 0.64 (Figure S4B). The regression slope of predicted on observed age was 0.42 with an intercept of 3.18 years. Mean absolute error was 3.35 years, and the median absolute error (MAE) was 2.79 years. Three individuals exhibited absolute prediction errors exceeding 6 years, including one exceeding 10 years, with a maximum absolute error of 15.14 years.

#### Bone ageing models fitted to combined sexes

*i) polynomial regression*

The pooled polynomial bone ageing model (combined sexes) yielded a Spearman correlation of ρ = 0.94 and an $R^2$ of 0.87 from the regression of predicted on observed age (Figure S5A). The regression slope was 0.88 with an intercept of 0.92 years. The mean absolute error was 1.75 years (SD = 1.42 years), with a median absolute error (MAE) of 1.32 years. Two individuals exhibited absolute prediction errors exceeding 6 years, and no individuals exceeded 10 years. The maximum absolute error was 6.45 years. The formula for the pooled polynomial regression is given as follows, where y denotes predicted age (years) and x the total bone score:

$$y\ (pooled) = -0.969052 + 0.0687288\,x + 0.0011129\ x^2$$

*ii) CAP model*

For the pooled CAP model, the first canonical axis (CAP1) showed a canonical correlation of δ = 0.91, corresponding to a squared canonical correlation ($\delta_1^2$) of 0.83. CAP-derived fitted values for individuals of known ages showed a mean absolute error of 1.90 years (median = 1.56 years; SD = 1.57 years), with a Spearman correlation of ρ = 0.92 (Fig. S5B). Linear regression yielded a slope of 0.88, an intercept of 0.84, and an $R^2$ of 0.85. Two individuals exhibited absolute errors larger than 6 years, with a maximum absolute error of 8.21 years.

### Text S3 Extended discussion

#### Sex-specific refitting of epigenetic clock published by Hanninger et al. (2005)

To ensure a valid comparison between bone ageing and epigenetic ageing, we refitted the epigenetic clock published by Hanninger et al. (2025) independently for females and males. Refitting the model by sex did not improve predictive performance. Correlation coefficients declined relative to the pooled model (MAE = 1.80, $r$ = 0.91, $R^2$ = 0.82; Hanninger et al., 2025), and mean absolute error increased in both sexes (females: MAE = 2.31, $r$ = 0.89, $R^2$ = 0.80; males: MAE = 2.79, $r$ = 0.80, $R^2$ = 0.64). These findings indicate that sex-specific refitting did not enhance age prediction accuracy within this population. The absence of improvement is consistent with prior evaluations of chronological DNA methylation clocks in humans, where sex-specific stratification has not consistently resulted in improved predictive performance (Carlsen et al., 2023; Sala et al., 2024).

In our modelling framework, epigenetic clock development required a training sample of approximately 70 individuals to achieve stable predictive performance (Mayne et al., 2021). Both sex-specific subsets fell well below this threshold. High-dimensional penalized regression approaches, such as elastic net, are sensitive to reductions in training size when the number of predictors greatly exceeds the number of samples, and the reduced sample sizes under stratification likely contributed to the observed decline in model performance (Mayne et al., 2021). Taken together, these results demonstrate that sex-specific refitting does not improve model performance within this study population. Accordingly, there is no empirical support for stratifying the epigenetic clock by sex under the present sample sizes.

#### Polynomial bone ageing model fitted to combined sexes

When fitted to the combined dataset, the polynomial bone ageing model demonstrated stronger predictive performance than the pooled epigenetic clock. The skeletal model achieved a Spearman correlation of $\rho = 0.94$ and an $R^2$ of 0.87, with a median absolute error (MAE) of 1.32 years. The CAP model achieved similar results with a MAE of 1.56 years, a Spearman correlation of $\rho = 0.92$ and an $R^2$ of 0.85. In comparison, the pooled epigenetic model yielded an MAE of 1.80 years, $r = 0.91$, and $R^2 = 0.82$ (Hanninger et al., 2025). Absolute prediction error was therefore lower in the skeletal model, and extreme errors were less frequent. These results indicate that, within this study population and under cross-validated conditions, radiographic bone maturation provides more precise chronological age estimates than the current methylation-based approach.

#### Supplementary tables

**Table S1:** Comparison of polynomial regression model performance for female common dolphins (*Delphinus delphis*) for using the total bone score calculated as the sum across all bone locations; differences between methods relate to the handling of the bones '*D2P2*' and '*D3P2*': *separate epiphyses* — proximal and distal epiphyses of these bones was scored separately; *combined score (average)* — each bone represented by the mean of proximal and distal epiphysis scores; c*ombined score (max)* — each bone represented by the higher of its proximal and distal epiphysis scores; *combined score (min)* — each bone represented by the lower of its proximal and distal epiphysis scores. The table reports absolute error distributions (SD of AE, Mean AE, Median AE) and predictive strength (Spearman correlation ρ) for each model. Due to bilateral asymmetry between left and right flipper, models were generated for the lower ('BSL'), higher ('BSH') and average bone score ('ABS') of both pectoral flippers, as well as the left ('Left') and right flipper ('Right') individually. Values are presented for model calibrated using the full dataset.

| | | | Separate epiphyses | | |
|---|---|---|---|---|---|
| **Metric** | **BSL** | **BSH** | **ABS** | **Left** | **Right** |
| SD of AE | 2.304426503 | 2.31692293 | 2.309711131 | 2.318220879 | 2.303496597 |
| Mean AE | 2.056128628 | 2.03443985 | 2.045139225 | 2.031725173 | 2.058469638 |
| Median AE | 1.384441195 | 1.392257652 | 1.380808347 | 1.393343285 | 1.37950293 |
| Correlation ρ | 0.912009911 | 0.911563645 | 0.911563645 | 0.911494831 | 0.911426016 |
| | | | **Combined score (average)** | | |
| **Metric** | **BSL** | **BSH** | **ABS** | **Left** | **Right** |
| SD of AE | 2.27222026 | 2.285164404 | 2.277467568 | 2.286876297 | 2.270586275 |
| Mean AE | 2.02850359 | 2.005279078 | 2.01679937 | 2.002029134 | 2.031667297 |
| Median AE | 1.38475681 | 1.35634075 | 1.354663124 | 1.369583993 | 1.382807018 |
| Correlation ρ | 0.912654231 | 0.912700107 | 0.912791859 | 0.912700107 | 0.912080783 |

**Table S1:** Comparison of polynomial regression model performance for female common dolphins (*Delphinus delphis*), continued.

| Combined score (max) | | | | | |
|---|---|---|---|---|---|
| **Metric** | **BSL** | **BSH** | **ABS** | **Left** | **Right** |
| SD of AE | 2.273700499 | 2.28801265 | 2.279550371 | 2.289931714 | 2.271852855 |
| Mean AE | 2.024175081 | 1.999372273 | 2.011617007 | 1.995726769 | 2.027714 |
| Median AE | 1.384114947 | 1.331029115 | 1.350514552 | 1.375987718 | 1.360803092 |
| Correlation ρ | 0.913352824 | 0.913077566 | 0.913742773 | 0.9133987 | 0.912458235 |
| **Combined score (min)** | | | | | |
| **Metric** | **BSL** | **BSH** | **ABS** | **Left** | **Right** |
| SD of AE | 2.271221478 | 2.282767565 | 2.275866168 | 2.28430509 | 2.269812542 |
| Mean AE | 2.032709613 | 2.011138276 | 2.021863857 | 2.008233523 | 2.035499785 |
| Median AE | 1.35122163 | 1.327043601 | 1.331230933 | 1.343365293 | 1.350051885 |
| Correlation ρ | 0.909741114 | 0.910268687 | 0.910268687 | 0.910268687 | 0.909741114 |

**Table S2:** Comparison of polynomial regression model performance for male common dolphins (*Delphinus delphis*) for using the total bone score calculated as the sum across all bone locations; differences between methods relate to the handling of the bones '*D2P2*' and '*D3P2*': *separate epiphyses* — proximal and distal epiphyses of these bones was scored separately; *combined score (average)* — each bone represented by the mean of proximal and distal epiphysis scores; c*ombined score (max)* — each bone represented by the higher of its proximal and distal epiphysis scores; *combined score (min)* — each bone represented by the lower of its proximal and distal epiphysis scores. The table reports absolute error distributions (SD of AE, Mean AE, Median AE) and predictive strength (Spearman correlation ρ). Due to bilateral asymmetry between left and right flipper, models were generated for the lower ('BSL'), higher ('BSH') and average bone score ('ABS') of both pectoral flippers, as well as the left ('Left') and right flipper ('Right') individually. Values are presented for model calibrated using the full dataset.

| | | Separate epiphyses | | | |
|---|---|---|---|---|---|
| **Metric** | **BSL** | **BSH** | **ABS** | **Left** | **Right** |
| SD of AE | 2.245175653 | 2.193492687 | 2.217891651 | 2.245561395 | 2.192899137 |
| Mean AE | 2.140668162 | 2.111756121 | 2.126465922 | 2.14028755 | 2.11230438 |
| Median AE | 1.404106133 | 1.387175281 | 1.381826617 | 1.399306545 | 1.400619043 |
| Correlation ρ | 0.941810777 | 0.943384396 | 0.942358123 | 0.942734423 | 0.941502895 |
| | | **Combined score (average)** | | | |
| **Metric** | **BSL** | **BSH** | **ABS** | **Left** | **Right** |
| SD of AE | 2.255090054 | 2.213887874 | 2.23349049 | 2.25684191 | 2.212126767 |
| Mean AE | 2.156476639 | 2.132520864 | 2.144821647 | 2.154219884 | 2.134741974 |
| Median AE | 1.395988273 | 1.407394249 | 1.38857795 | 1.364783817 | 1.401119946 |
| Correlation ρ | 0.941023968 | 0.942118659 | 0.942221286 | 0.942289704 | 0.941331849 |
| | | **Combined score (max)** | | | |
| **Metric** | **BSL** | **BSH** | **ABS** | **Left** | **Right** |
| SD of AE | 2.255507061 | 2.209047971 | 2.230968893 | 2.257536542 | 2.20699857 |
| Mean AE | 2.152772143 | 2.124930736 | 2.139269899 | 2.149945008 | 2.127724889 |
| Median AE | 1.395501583 | 1.393517868 | 1.374373473 | 1.352211772 | 1.398486689 |
| Correlation ρ | 0.942358123 | 0.943863323 | 0.943555441 | 0.943623859 | 0.942118659 |

**Table S2:** Comparison of polynomial regression model performance for male common dolphins (*Delphinus delphis*), continued

| | Combined score (min) | | | | |
|---|---|---|---|---|---|
| **Metric** | **BSL** | **BSH** | **ABS** | **Left** | **Right** |
| SD of AE | 2.254670811 | 2.219025591 | 2.236184689 | 2.256193361 | 2.217195708 |
| Mean AE | 2.160316464 | 2.140031988 | 2.150337253 | 2.158573378 | 2.142051411 |
| Median AE | 1.385035199 | 1.421344065 | 1.398018656 | 1.377086 | 1.40268362 |
| Correlation ρ | 0.940750295 | 0.941844986 | 0.941810777 | 0.942016031 | 0.941058177 |

**Table S3:** Polynomial regression models describing the relationship between total pectoral flipper bone score and chronological age in common dolphins (*Delphinus delphis*). For each model, two aspects vary: (1) the treatment of the second phalanges of digits II and III ('D2P2' and 'D3P2'), and (2) the handling of bilateral differences between left and right flippers. With respect to 'D2P2' and 'D3P2', four scoring approaches are shown: separate epiphyses, where the proximal and distal epiphyses are scored independently and each epiphysis contributes separately to the total score; combined score (average), where each bone is represented by the mean of its proximal and distal epiphysis scores; combined score (max), where each bone is represented by the higher of its two epiphysis scores; and combined score (min), where each bone is represented by the lower of its two epiphysis scores. Bilateral asymmetry is addressed using five variants: 'BSL', based on the lower-scoring flipper; 'BSH', based on the higher-scoring flipper; 'ABS', based on the average of left and right flipper scores; and separate models using the left ('Left') and right ('Right') flippers individually. The table lists the resulting second-order polynomial equations of the form $y = \beta_0 + \beta_1 x + \beta_2 x^2$, where x is the total bone score and y is predicted age (years). These equations were calculated based on the full data set.

| Model | Females | Males |
|---|---|---|
| | Separate epiphyses | |
| 'BSL' | $y = -0.7376 + 0.036396 \times x + 0.00117899 \times x^2$ | $y = 0.158557 + 0.0271375 \times x + 0.00105291 \times x^2$ |
| 'BSH' | $y = -0.829114 + 0.0379345 \times x + 0.00117051 \times x^2$ | $y = 0.146565 + 0.0268819 \times x + 0.00105153 \times x^2$ |
| 'ABS' | $y = -0.780987 + 0.0370384 \times x + 0.0011759 \times x^2$ | $y = 0.155889 + 0.02681 \times x + 0.00105394 \times x^2$ |
| 'Left' | $y = -0.761154 + 0.0361074 \times x + 0.00118189 \times x^2$ | $y = 0.121697 + 0.0279779 \times x + 0.0010484 \times x^2$ |
| 'Right' | $y = -0.801872 + 0.0380927 \times x + 0.00116854 \times x^2$ | $y = 0.182992 + 0.0260534 \times x + 0.00105596 \times x^2$ |
| | Combined score average | |
| 'BSL' | $y = -1.00156 + 0.0471288 \times x + 0.00144368 \times x^2$ | $y = 0.0528455 + 0.0334955 \times x + 0.00130271 \times x^2$ |
| 'BSH' | $y = -1.07108 + 0.0480113 \times x + 0.00143955 \times x^2$ | $y = 0.0507911 + 0.0328258 \times x + 0.00130549 \times x^2$ |
| 'ABS' | $y = -1.03375 + 0.0474111 \times x + 0.00144324 \times x^2$ | $y = 0.053803 + 0.0330274 \times x + 0.00130539 \times x^2$ |
| 'Left' | $y = -0.99657 + 0.04576 \times x + 0.00145529 \times x^2$ | $y = 0.0223684 + 0.0343353 \times x + 0.0012974 \times x^2$ |
| 'Right' | $y = -1.07363 + 0.0492808 \times x + 0.00142874 \times x^2$ | $y = 0.0808683 + 0.0319989 \times x + 0.0013107 \times x^2$ |

**Table S3:** Polynomial regression models describing the relationship between total pectoral flipper bone score and chronological age in common dolphins (*Delphinus delphis*), continued.

| Model | Females | Males |
|---|---|---|
| | Combined score (max) | |
| 'BSL' | $y = -1.03169 + 0.0474985 \times x + 0.00144261 \times x^2$ | $y = 0.0451948 + 0.0331951 \times x + 0.00130663 \times x^2$ |
| 'BSH' | $y = -1.08317 + 0.0476272 \times x + 0.00144443 \times x^2$ | $y = 0.0444677 + 0.0325156 \times x + 0.00130866 \times x^2$ |
| 'ABS' | $y = -1.05451 + 0.0473807 \times x + 0.00144539 \times x^2$ | $y = 0.0475891 + 0.0326732 \times x + 0.00130939 \times x^2$ |
| 'Left' | $y = -1.01363 + 0.0455433 \times x + 0.0014589 \times x^2$ | $y = 0.0157955 + 0.0340519 \times x + 0.00130103 \times x^2$ |
| 'Right' | $y = -1.09958 + 0.0495122 \times x + 0.0014589 \times x^2$ | $y = 0.0735636 + 0.0316692 \times x + 0.00131418 \times x^2$ |
| | Combined score (min) | |
| 'BSL' | $y = -0.970026 + 0.0467022 \times x + 0.00144517 \times x^2$ | $y = 0.0613129 + 0.0337571 \times x + 0.00129908 \times x^2$ |
| 'BSH' | $y = -1.05726 + 0.0483327 \times x + 0.0014351 \times x^2$ | $y = 0.0574401 + 0.0331287 \times x + 0.00130233 \times x^2$ |
| 'ABS' | $y = -1.0115 + 0.0473849 \times x + 0.0014415 \times x^2$ | $y = 0.0606674 + 0.0333535 \times x + 0.00130158 \times x^2$ |
| 'Left' | $y = -0.977924 + 0.0459177 \times x + 0.00145206 \times x^2$ | $y = 0.0297298 + 0.034582 \times x + 0.00129405 \times x^2$ |
| 'Right' | $y = -1.04608 + 0.0489863 \times x + 0.00142926 \times x^2$ | $y = 0.0885223 + 0.0323195 \times x + 0.00130724 \times x^2$ |

#### Supplementary figures

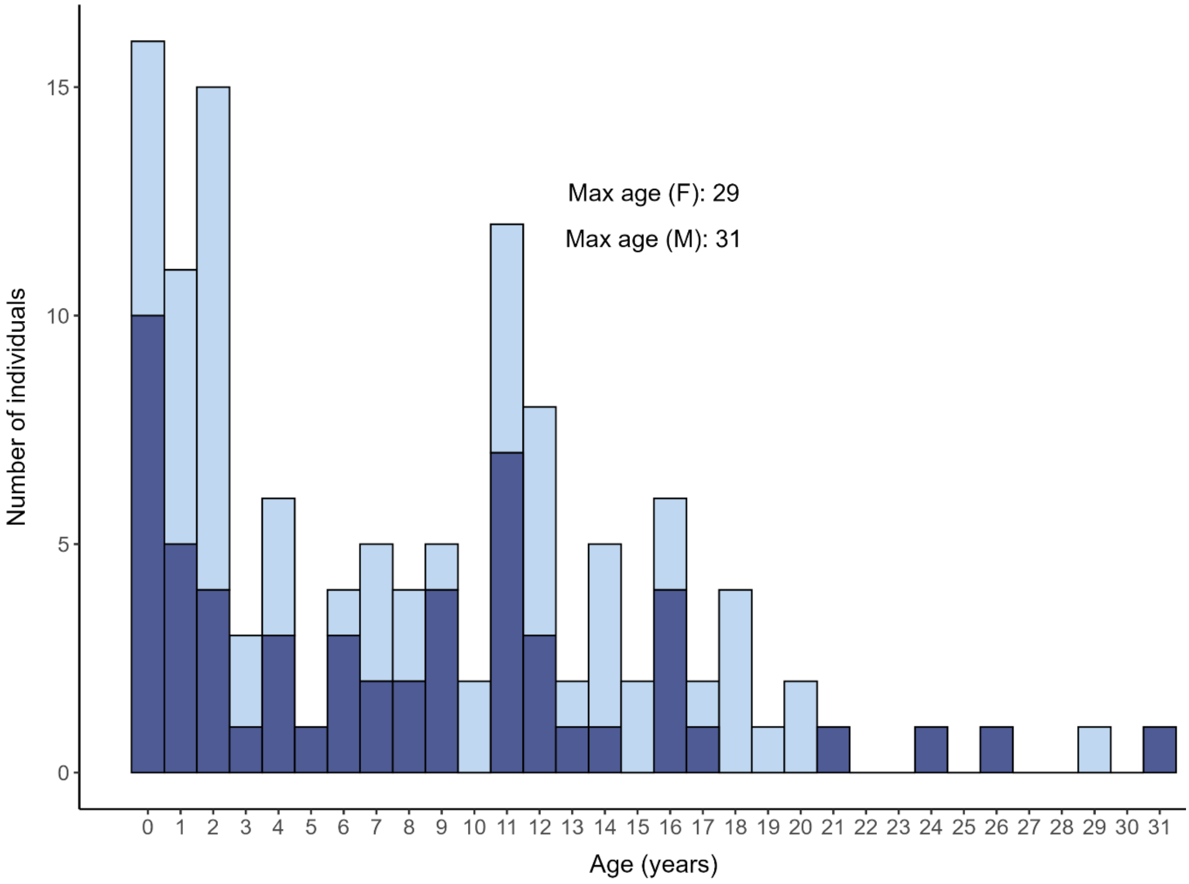


**Figure S1**: Dental age distributions by sex for common dolphins (*Delphinus delphis*) for which pectoral flipper bone maturation was assessed. The figure does not include foetuses.

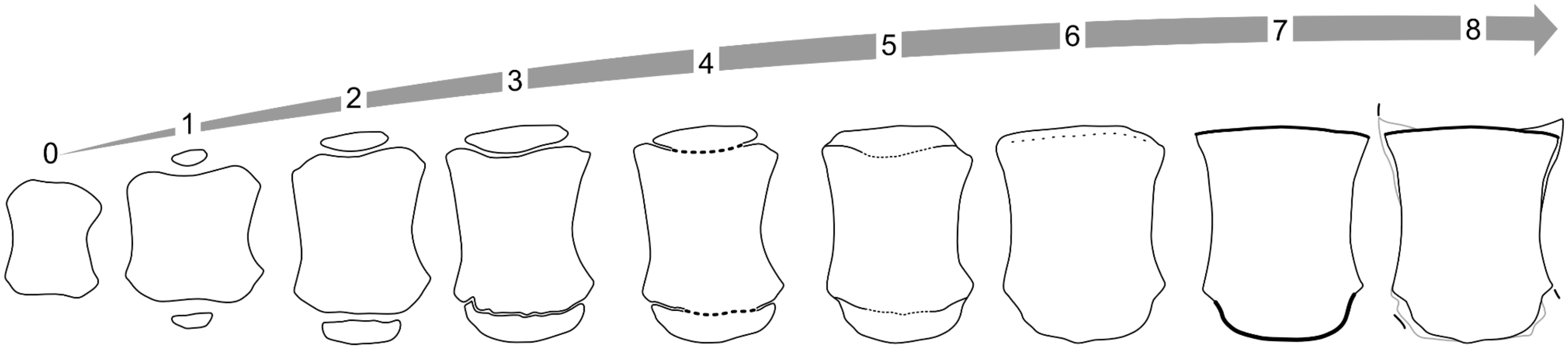


**Figure S2.** Long bone scoring scheme, reprinted from (Barratclough et al., 2019) and illustrated by Veronica Candejas: **Score -1 (not shown):** No primary ossification centre visible. **Score 0:** The primary centre is present, but no secondary ossification centre is visible. **Score 1:** An epiphyseal ossification centre is present but only occupies <50% of the width of the adjacent metaphysis. **Score 2:** The secondary ossification centre is well established, ranging between 50–100% of the metaphyseal width. The physis is evident as a distinct radiolucent space, while the secondary centre appears less mature and irregular in depth across its width. Maturation progresses in a midline-to-abaxial direction. **Score 3:** Thinning of the radiolucent physis but no evidence of fusion of the metaphysis and the secondary ossification centre. Mineral density increases in the subchondral bone on both sides of the physis. Late stage 2 can resemble early stage 3; the distinction lies in whether physis width is irregular (stage 2) or uniform (stage 3). **Score 4:** Osseous bridges begin to form; in the radius and ulna this starts midline and proceeds abaxially. The abaxial margins remain open, and both early bridge formation and >90% bridging qualify as stage 4. **Score 5:** Complete closure is present, accompanied by a faint hyperdense "ghost" physeal line or epiphyseal scar spanning at least 50% of the plate. The former epiphysis may not reach the full transverse width of the bone, but closure is complete. **Score 6:** The physeal line is remodelled, with <50% to no evidence of the hyperminerlized transverse remnant, and the former epiphysis extends across the full transverse width of the bone. **Score 7:** Complete fusion occurs with no evidence of a previous ghost physeal plate and flattening of the physeal surfaces with accentuated pointed edges. **Score 8:** Arthritic changes are visible, including osteophytes, osteolysis, articular calcification, and in some cases fusion of metacarpals or phalanges.

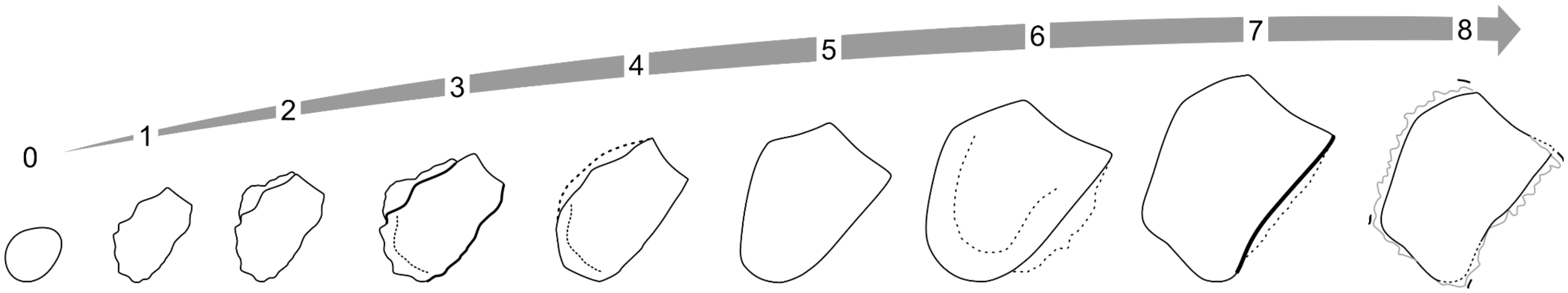


**Figure S3.** Delta bone scoring scheme, reprinted from (Barratclough et al., 2019) and illustrated by Veronica Candejas: **Stage -1** (not shown): Not present. **Stage 0:** Very small oval shape, undefined delta shape. **Stage 1:** Axial surface starts to be linear, consistent with delta shape. **Stage 2:** One flat surface. Areas of slight irregular perimetral surface typically on the proximal side with a degree of adjacent mineralization which can oppose the epiphyseal surface but not necessarily show definitive osseous bridge formation. Bone density is clearly reduced in peripheral new bone in comparison to the primary ossification centre. **Stage 3:** Increased mineralization of adjacent proximal cartilaginous surface, consolidation may not be present. **Stage 4:** Area increases in consolidation and mineralization, slightly less dense than the body of the delta bone. A defined hyperminerlized physeal plate line is present. The abaxial aspect can show the initiation of the new straight lateral border rather than a horseshoe shape. **Stage 5:** Density of the secondary ossification centre is similar to primary bone and initial mineralization is present on the distal surface, with the proximal side showing a defined straight lateral border rather than a horseshoe shape. Individual variation may result in non-linear borders; therefore, emphasis should be placed on the secondary ossification centres on the proximal and distal surfaces. **Stage 6:** Smooth proximal surface with matching density to the primary ossification centre, distal surface still shows some incongruent mineralization. Physeal line is still visible, denoted by a dotted line. **Stage 7:** Smooth proximal surface with complete consolidation of the secondary ossification centre on both sides and dissolution of the physeal line. **Stage 8:** Degenerative changes present.

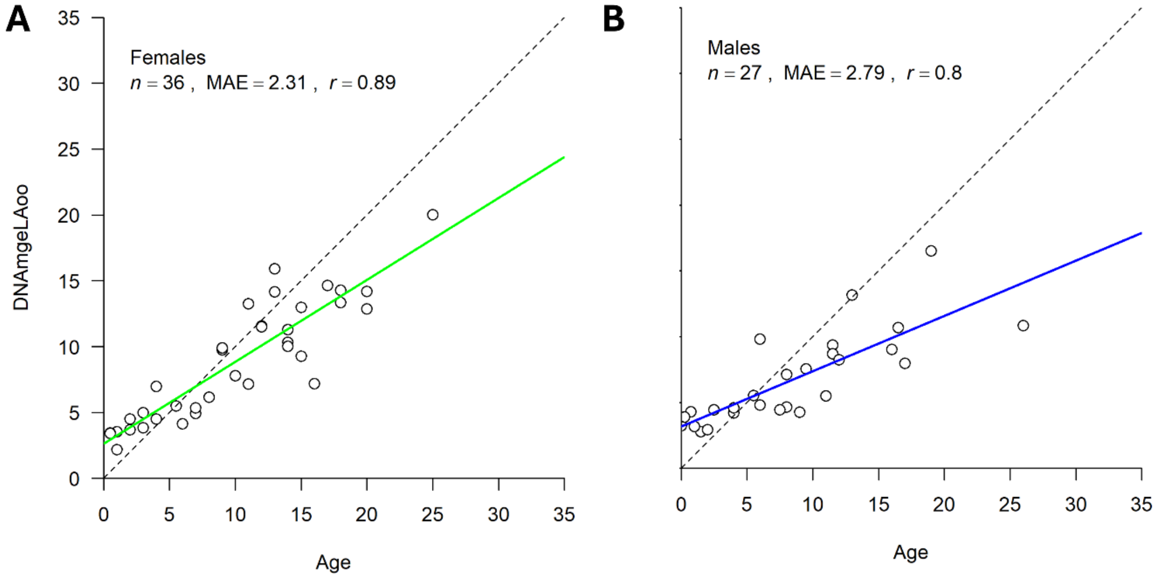


**Figure S4:** Sex-specific refitting of the epigenetic age model originally published by Hanninger et al. (2025) using the restricted subset of common dolphins (*Delphinus delphis*). The model was refitted separately for **A)** females and **B)** males to evaluate whether sex-specific stratification improves predictive performance relative to the pooled model. DNAmAgeLOO (leave-one-out cross-validated predictions) is plotted against chronological age. Solid lines represent linear regression fits; dashed lines indicate the 1:1 relationship. Sample size (*n*), mean absolute error (MAE), and Pearson correlation coefficient (*r*) are shown for each sex.

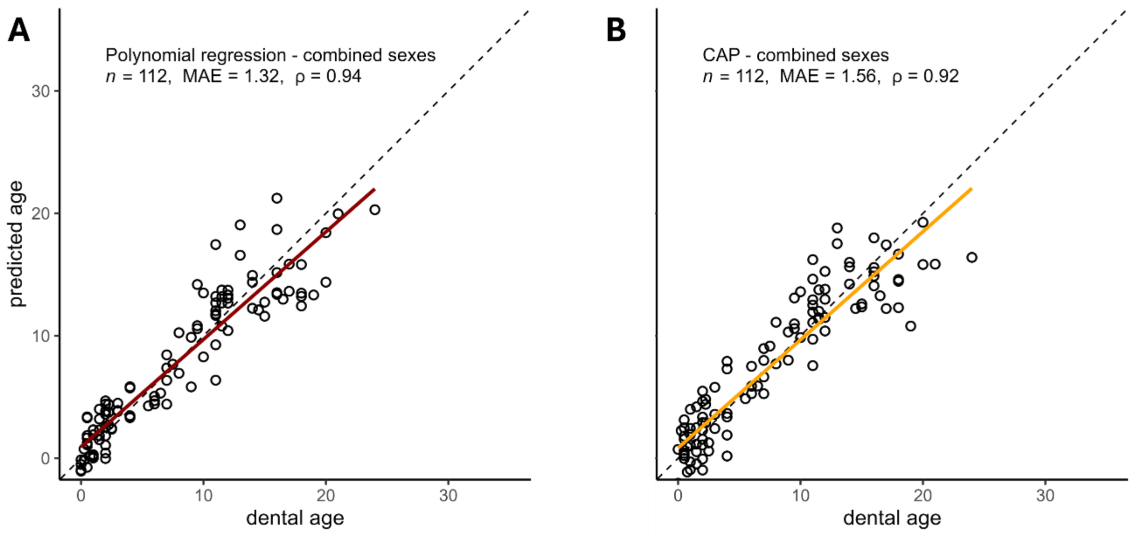


**Figure S5:** Relationship between dental age and predicted age for common dolphins (*Delphinus delphis*). **A)** Pooled polynomial bone-ageing model (combined sexes; outliers excluded). Predicted ages were generated via leave-one-out cross-validation (LOOCV) of a second-degree polynomial regression fitted to ossification scores derived from radiographic assessment of the right pectoral flipper. **B)** LOOCV-predicted ages from the CAP model. The solid red (polynomial regression) and orange (CAP) lines represent linear regressions of predicted on observed age, and the dashed line indicates the 1:1 relationship. Model performance metrics (sample size, *n*; median absolute error, MAE; and Spearman's rank correlation, ρ) are provided within each panel.
